# Cystinosin regulates kidney inflammation through its interaction with galectin-3

**DOI:** 10.1101/289702

**Authors:** Tatiana Lobry, Roy Miller, Nathalie Nevo, Celine J. Rocca, Jinzhong Zhang, Sergio D. Catz, Fiona Moore, Anne Bailleux, Ida Chiara Guerrera, Marie-Claire Gubler, Cheung W. Wilson, Robert H. Mak, Tristan Montier, Corinne Antignac, Stephanie Cherqui

## Abstract

Inflammation is implicated in the pathogenesis of many disorders. Here, we show that cystinosin, protein defective in the lysosomal storage disorder cystinosis, is a critical regulator of galectin-3 during inflammation. Cystinosis is a lysosomal storage disorder and despite ubiquitous expression of cystinosin, kidney is the primary organ to be impacted by the disease. Here, we show that cystinosin interacts with galectin-3 and enhances its lysosomal localization and degradation. Galectin-3 is also found overexpressed in the kidney of the mouse model of cystinosis, *Ctns^-/-^*mice. Absence of galectin-3 in *Ctns^-/-^* mice led to a better renal function and structure, and decreased macrophage/monocyte infiltration in the kidney. Finally, galectin-3 interacts with a protein implicated in the recruitment of monocytes and macrophages during inflammation, Monocyte Chemoattractant Protein-1 (MCP1), that was found increased in the serum of *Ctns^-/-^* mice. These findings highlight a new role of cystinosin and galectin-3 interaction in inflammation, providing a mechanistic explanation for kidney disease pathogenesis in cystinosis, which may lead to the identification of new drug targets to delay its progression.

## Introduction

Inflammation is a normal acute response of the organism to injury or infection (Medzhitov, 2008), but chronic inflammation is associated with tissue damage. In the case of chronic disease, such as diabetes or obesity-related, damaged tissue activate the immune system, which lead to continuous activation of inflammation and thus, to tissue degeneration (Donath, 2013). Cytokines are key modulators of inflammation capable of activating or resolving the inflammation process (Jaffer, Wade et al., 2010). In patients with Chronic Kidney Disease (CKD), elevated plasma concentrations of cytokines (IL-6 and TNFa) and inflammation markers (CRP, IL-6, hyaluronan and neopterin) are associated with progression to End-Stage Renal Disease (ESRD) (Pecoits-Filho, Heimburger et al., 2003). Understanding the specific mediators that trigger inflammatory responses would allow the discovery of appropriate drug target to prevent degenerative damage.

Cystinosis is an autosomal recessive metabolic disease that belongs to the family of the lysosomal storage disorders, characterized by the accumulation of cystine within all organs (Gahl, Thoene et al., 2002). The gene defective in cystinosis, *CTNS,* encodes the lysosomal cystine/proton co-transporter, cystinosin (Cherqui, Kalatzis et al., 2001, Kalatzis, Cherqui et al., 2001). Although *CTNS* is ubiquitously expressed, kidney is the first tissue impacted in cystinosis. Patients typically present in their first year of life with symptoms of Fanconi syndrome characterized by severe fluid and electrolyte disturbances, poor growth and rickets (Cherqui & Courtoy, 2016). Patients subsequently develop chronic kidney disease that progresses to ESRD requiring renal replacement therapy. Cystine accumulation in all tissues eventually leads to multi-organ dysfunction and patients suffer from photophobia and blindness, hypothyroidism, hypogonadism, diabetes, myopathy, and central nervous system deterioration (Nesterova & Gahl, 2008). In the appropriate background, a knockout mouse model (*Ctns^-/-^* mice) (Cherqui, Sevin et al., 2002) largely replicates the kidney phenotype of cystinotic patients (Nevo, Chol et al., 2010) as well as deposition of cystine crystals in the cornea (Kalatzis, Serratrice et al., 2007) and thyroid dysfunction (Gaide Chevronnay, Janssens et al., 2015).

Besides supportive therapy, the current treatment for cystinosis is the substrate depletion drug cysteamine, which delays the progression of cystinosis complications but has no effect on the renal Fanconi syndrome and does not prevent ESRD (Brodin-Sartorius, Tete et al., 2012, Cherqui, 2012, Gahl, Balog et al., 2007). The specific sensitivity of the kidneys in cystinosis is still not fully understood as cystine accumulates in all tissues (Cherqui & Courtoy, 2017). Recently, new cellular pathways have been shown to be impacted by the absence of cystinosin beyond cystine transport. As such, chaperone mediated autophagy (CMA) is impaired and LAMP2a, main protagonist of CMA, is mislocalized in murine *Ctns*-deficient cells and tissues (Napolitano, Johnson et al., 2015a). Physical interaction of cystinosin with almost all components of v-ATPase, ragulator and RagA/RagC was recently demonstrated as well as defective lysosomal recruitment of mTOR upon nutrient shortage (Andrzejewska, Nevo et al., 2016). TFEB expression, regulator of lysosomal clearance, autophagy and anabolism, was also recently shown to be impaired in cystinosin-deficient proximal tubular cells (Rega, Polishchuk et al., 2016). A striking common property of these studies is that defects in mTOR signaling, TFEB expression and CMA could not be corrected upon full cystine clearance by cysteamine, showing that these cellular anomalies are due to the very absence of cystinosin, beside cystine overload (Andrzejewska et al., 2016, Napolitano et al., 2015a, Rega et al., 2016). In the present study, we reveal a new role of cystinosin in inflammation through its interaction with galectin-3 (Gal3), a member of the lectin and beta-galactoside-binding protein family (Dumic, Dabelic et al., 2006). We provide evidence that, in cystinosis, absence of cystinosis leads to Gal3 overexpression in kidneys enhancing macrophage infiltration and chronic kidney disease progression. In addition, we identify Monocyte Chemoattractant Protein-1 (MCP1) as a potential mediator, revealing new gene targets for drug therapy.

## Results

### Gal3 interacts with cystinosin via its carbohydrate recognition domain

To identify specific interaction partners using quantitative mass spectrometry, MDCK cells were transduced with a lentiviral construct to stably express the cystinosin-GFP fusion protein. In addition to the 9 subunits of the vATPase published previously (Andrzejewska et al., 2016), mass spectrometry analysis of anti-GFP immunoprecipitates identified Gal3 and galectin-9 (Figure 1A). This interaction of cystinosin and Gal3 was further confirmed by co-immunoprecipitation experiments with anti-GFP and anti-Gal3 antibodies followed by western blotting (Figure 1B and 1C). In addition, we showed that interaction between Gal3 and cystinosin was mediated by Gal3 Carbohydrate Recognition Domain (CRD) localized in the C-terminal tail of the protein. Indeed, interaction was inhibited by thiodigalactoside, a potent inhibitor of galectin-carbohydrate interactions, due to its high affinity for Gal3 (Figure 1D). Like all galectins, Gal3 has an affinity for glycosylated proteins (Dumic et al., 2006), so it is likely that the interaction with Gal3 occurs through the cystinosin glycosylated moieties located in the N-terminal tail of the protein localized within lysosome lumen (Nevo, Thomas et al., 2017). This, in turn, suggests that Gal3 can enter lysosomes.

**Figure 1:**
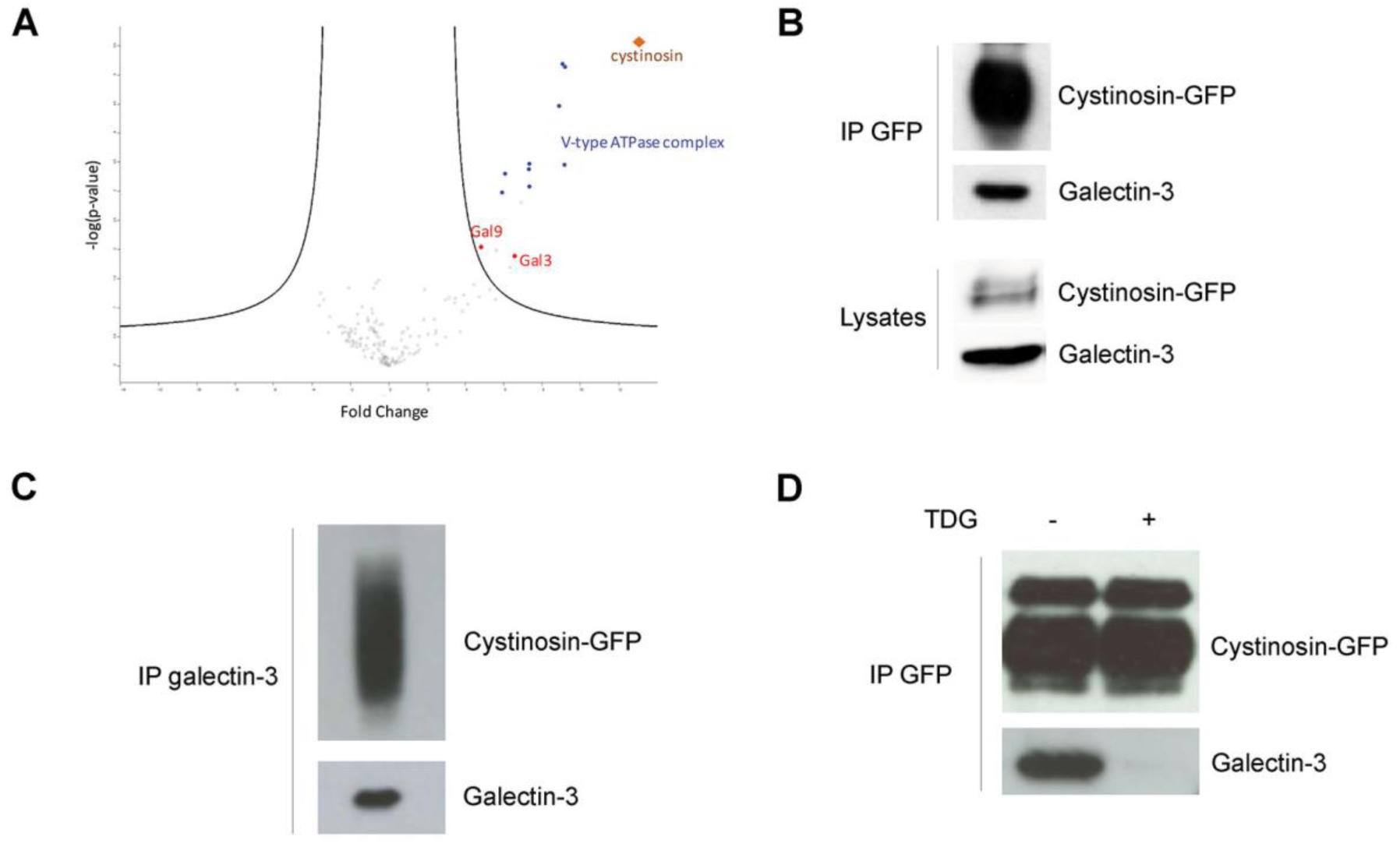
Gal3 interacts with cystinosin-GFP via its carbohydrate recognition domain. **a** Volcano plot representation of proteins immunoprecipitated by the anti-GFP antibody and identified by mass spectrometry (nanoRSLC-Q Exactive Plus MS) in MDCK cells overexpressing cystinosin-GFP vs non-transfected cells (4 independent experiments). Lysates of MDCK cells stably expressing cystinosin-GFP were immunoprecipitated with **b** anti-GFP or **c** anti-Gal3 antibodies, and coimmunoprecipitated proteins were analyzed by western blotting (n=3). IP, Immunoprecipitation. **d** Lysates of MDCK cells stably expressing cystinosin-GFP were treated or not for 30 minutes with 5mM thiodigalactoside (TDG), a potent inhibitor of galectin-carbohydrate interactions. Lysates were then immunoprecipitated with anti-GFP antibodies and coimmunoprecipitated proteins were analyzed by western blotting (n=3).

### Cystinosin enhances Gal3 lysosomal localization and degradation

Gal3 can localize in nucleus, cytoplasm or at the cell surface (Dumic et al., 2006). To verify the potential lysosomal localization of Gal3 when interacting with cystinosin, we treated cystinosin-GFP-expressing MDCK cell lysates with proteinase K in the presence or absence of 2% Triton X-100. Gal3, such as cathepsin D (an intralysosomal protease), was protected from digestion by proteinase K whereas both proteins were digested when lysosomes were permeabilized with Triton X-100 (Figure 2A). Similar data were obtained in lysosomes isolated from mouse liver, confirming lysosomal localization of Gal3 *in vivo* (Figure 2B and 2C). Immunostaining of cystinosin-GFP-expressing MDCK cell lines with anti-galectin-3 antibodies confirmed the presence of Gal3 in the lumen of the lysosomes while cystinosin-GFP and Lamp-2 (a lysosomal transmembrane protein) were found as expected only at the membrane of the vesicles (Figure 2D). However, this pattern was predominant only when MDCK overexpressed cystinosin, suggesting that cystinosin is involved in the lysosomal localization of Gal3.

**Figure 2:**
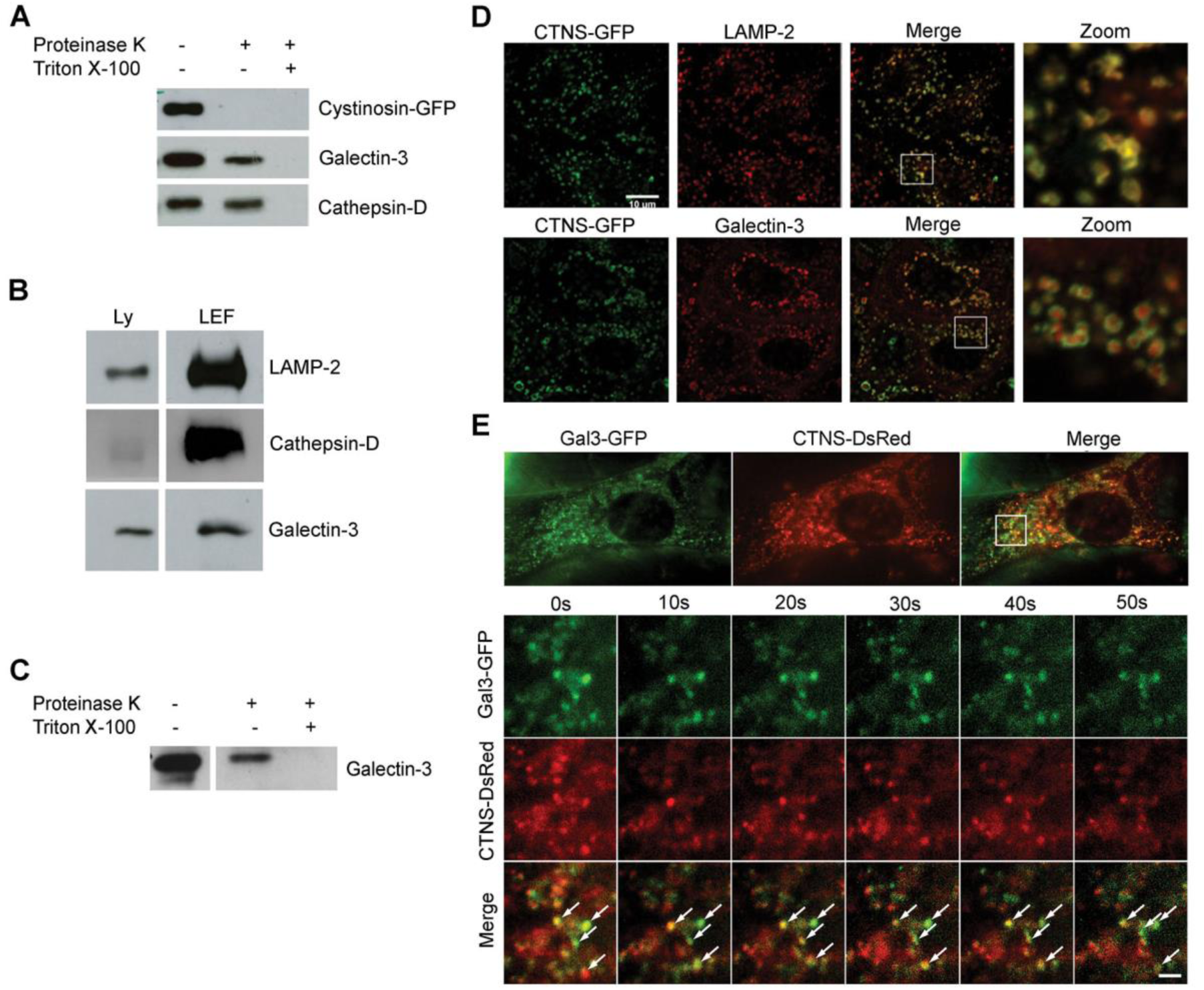
Gal3 is localized into the lumen of late endosomes and lysosomes. **a** Lysates of MDCK cells stably expressing cystinosin-GFP were treated or not with 2.5 µg/ml proteinase K for 30 min at 4°C, in the presence or in the absence of 2% Triton X-100 that permeabilized internal membranes. The samples were loaded on SDS-PAGE and processed for immunoblot with either anti-GFP (localized on the outer aspect of the lysosome), -cathepsin-D (intra-lysosomal hydrolase) or -Gal3 antibodies. **b** Lysates (Ly) or lysosome-enriched fractions (LEF) from mouse liver were probed with the indicated antibodies. **c** Lysosome-enriched fractions were treated or not with 25 µg/ml proteinase K for 30 min at 4°C, in the presence or in the absence of 2% Triton X-100. The samples were loaded on SDS-PAGE and processed for immunoblot with anti-gal3 antibodies. **d** Immunostaining of late endosomes and lysosomes with Lamp-2 in cystinosin-GFP MDCK cell lines revealed a co-localization between cystinosin-GFP and Lamp-2 at the lysosomal membrane. Gal3 is present within the vesicles whereas cystinosin-GFP is located at the membrane of the vesicles. Scale bar: 10 µm. **e** *Ctns^-/-^* MEFs expressing both Gal3-GFP and cystinosin-DsRed were analyzed by TIRFM. Vesicular trafficking was monitored during 60 seconds and analyzed using Image*J*. Magnifications of the selected areas are represented in the bottom panels and show true colocalization as determined by the spatiotemporal co-distribution of the proteins (indicated with arrows). A total of 9 cells were analyzed for each condition. Scale bar: 2 µm. The dynamics of the labeled vesicles and the spatiotemporal distribution of Gal3 and cystinosin can be viewed in associated Supplementary movies S1 and S2.

To investigate if cystinosin is an important component of the molecular machinery necessary for Gal3 trafficking, we simultaneously analyzed the dynamics of both cystinosin and Gal3-positive vesicles using dual-color live cell Total Internal Reflection Fluorescence microscopy (TIRFM) in mouse embryonic fibroblasts (MEF) from wild type (WT) and *Ctns^-/-^* mice expressing both CTNS-DsRed and Gal3-GFP. Our analysis confirms that cystinosin and Gal3 undergo true colocalization in *Ctns^-/-^* MEF as validated by the similar spatiotemporal distribution of these molecules in the TIRFM zone (Figure 2E). Data in WT MEF were similar and therefore, not shown. Moreover, MEF transfected only with Gal3-GFP were analyzed and Gal3-GFP was found everywhere in the cytoplasm of the cells and no distinct vesicular structure was observed for Gal3-GFP contrarily to the co-transfection with CTNS-DsRed (data not shown).

These data were confirmed in 293T cells, in which strong GFP signal in the cytoplasm was exclusively seen in cells expressing only Gal3-GFP whereas in cells overexpressing both Gal3-GFP and cystinosin-DsRed, Gal3-GFP was mainly localized within cystinosin-DsRed-expressing lysosomes (Figure 3A). Because the GFP signal was much lighter in the Gal3 and cystinosin co-transfected cells, suggesting the presence of less Gal3 protein, we quantified the presence of Gal3 in cells transfected with Gal3-GFP alone or Gal3-GFP and CTNS-DsRed, and Gal3-GFP and DsRed as control using western blot analysis. We found significantly less Gal3-GFP proteins in cells co-transfected with CTNS-DsRed compared to cells transfected with Gal3-GFP alone or co-transfected with DsRed (Figure 3B). Altogether, these results suggest that cystinosin enhances the lysosomal localization of Gal3 and degradation.

**Figure 3:**
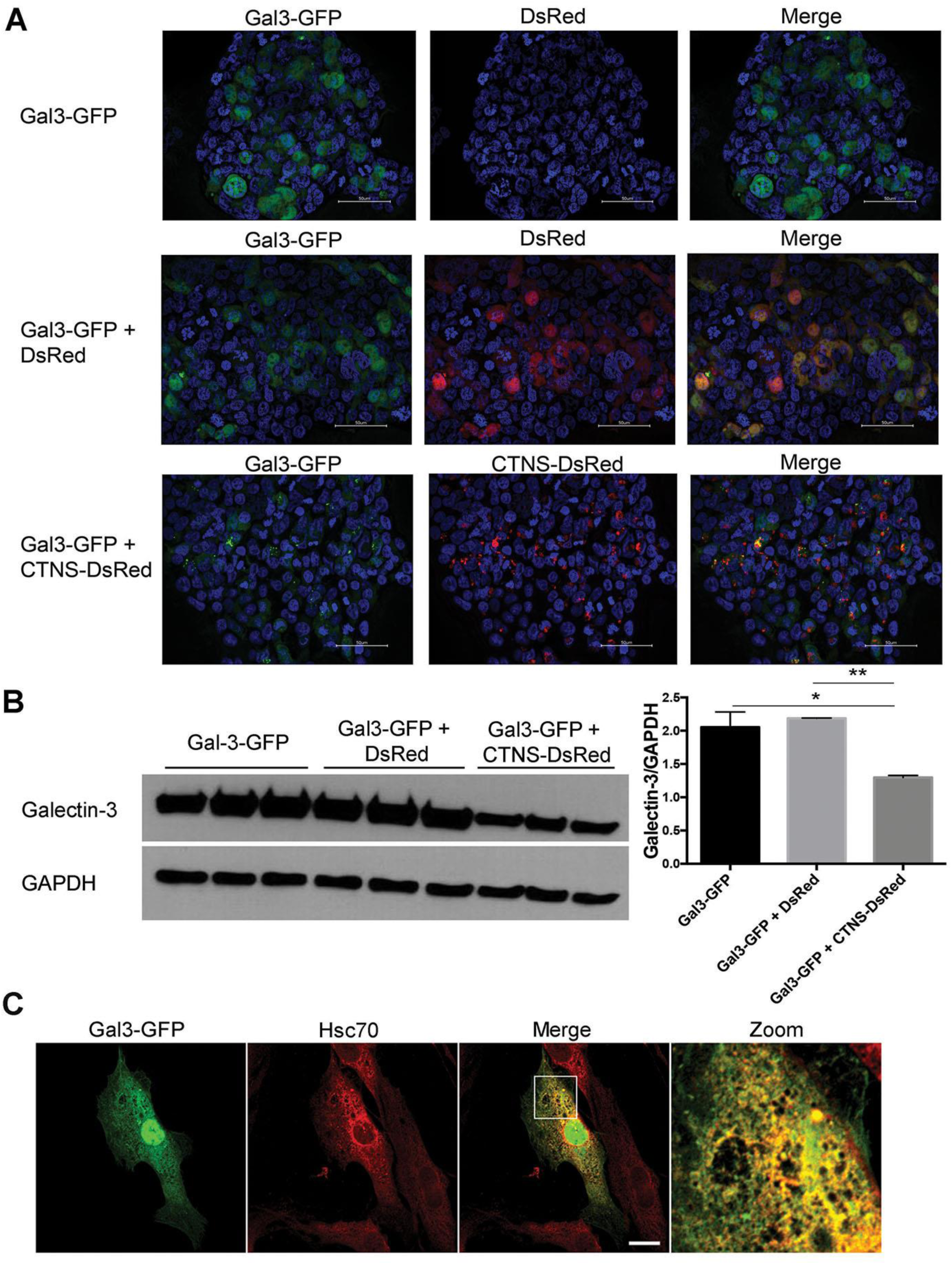
Cystinosin enhances Gal3 lysosomal localization and degradation. **a** Fluorescent images of large views of 293T cells transfected with Gal3-GFP alone or with either DsRed or cystinosin-DsRed. Scale bar: 50 µm. **b** Proteins were isolated from the transfected 293T cells, resolved on a SDS-PAGE, revealed with an anti-Gal3 or anti-GAPDH antibody as reference and quantified using Image J Software, n=3 and statistical test used was one-way ANOVA followed by Tukey’s test. *P<0.05; **P<0.01. **c** *Ctns^-/-^*MEFs were transfected with Gal3-GFP expression vector and immunostained for endogenous Hsc70. Scale bar: 20 µm.

Cystinosin has been previously shown to be involved in CMA (Napolitano, Johnson et al., 2015b). Heat shock 70 KDa protein (Hsc70) participates in CMA by aiding the unfolding and translocation of substrate proteins across the membrane into the lysosomal lumen in a LAMP2A-dependent manner (Majeski & Dice, 2004, Xie, Zhang et al., 2015). Because Gal3 was proposed to be degraded by CMA (Li, Ma et al., 2010), we investigated the localization of Gal3 related to that of Hsc70 in cystinotic cells. Confocal microscopy analysis confirmed the colocalization of Gal3-GFP at Hsc70-positive structures in both WT (data not shown) and *Ctns^-/-^* cells (Figure 3C), supporting the co-transportation of Gal3 with Hsc70, and suggesting that lysosomal internalization but not Gal3 recognition by the chaperone is defective in cystinosis.

### Gal3 is involved in kidney inflammation in *Ctns^-/-^*mice

To study the role of Gal3 in cystinosis *in vivo*, we generated a double knock-out mouse model deficient in both cystinosin and Gal3, (*Ctns^-/-^ Gal3^-/-^* mice). WT, *Ctns^-/-^*and *Gal3^-/-^* mice were used as controls. Cystine content was measured in the kidney of 8-9 months and 12-15 months old mice and because cystine content is different in male and female *Ctns^-/-^* kidneys, they were analyzed separately. No significant difference was observed in male and female kidneys from *Ctns^-/-^ Gal3^-/-^*mice compared to *Ctns^-/-^* mice at 8-9 months (data not shown) and 12-15 months of age (Figure 4A).

**Figure 4:**
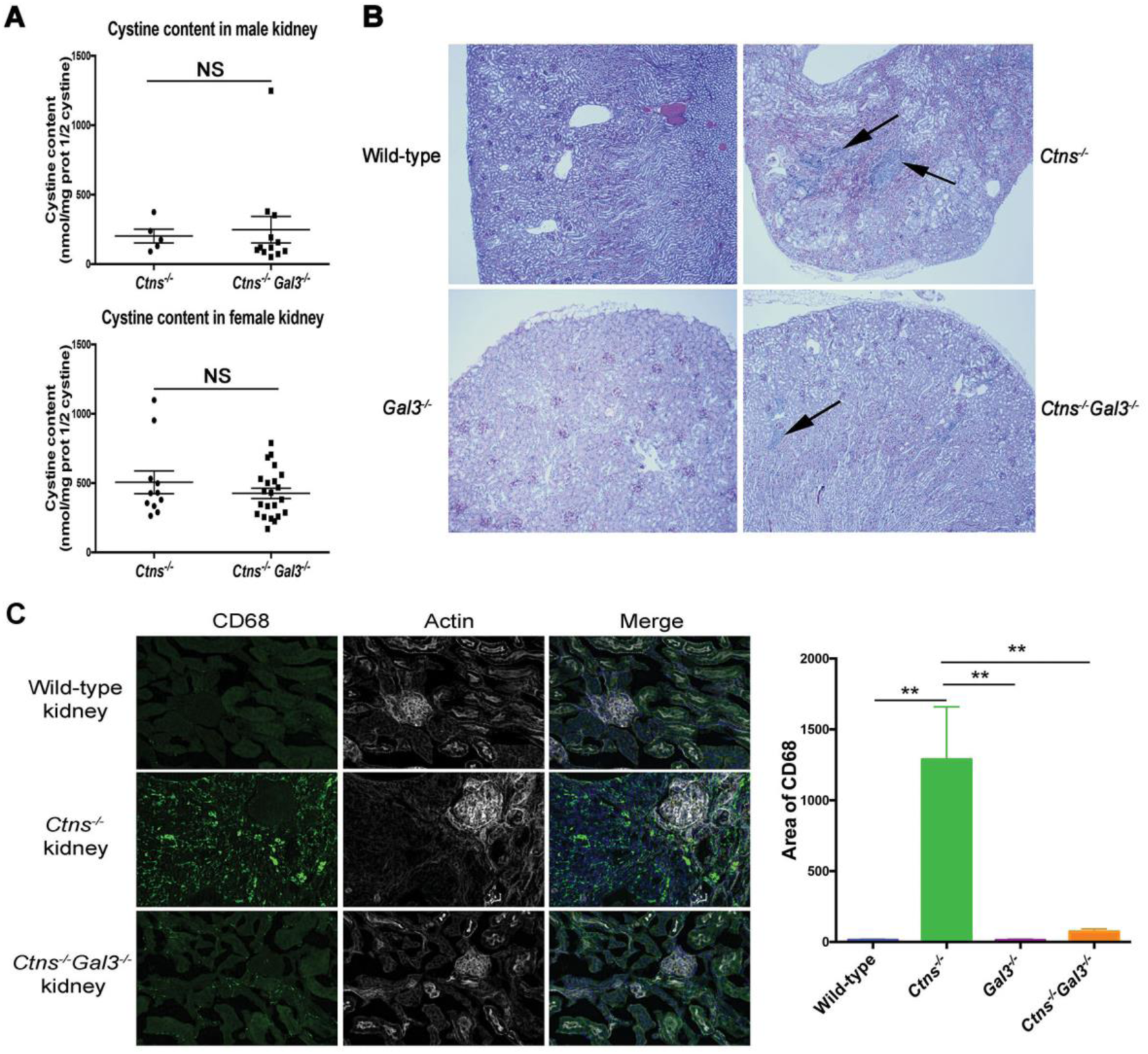
Effect of the absence of Gal3 on renal function and kidney structure in 12-15-month-old mice. **a** Cystine content levels (nmol half cystine/mg protein) in *Ctns^-/-^* (n=16, 5 males and 11 females) and *Ctns^-/-^ Gal3^-/-^* (n=34, 12 males and 22 females) at 12-15 months in the different tissues tested. *P* values were determined using unpaired 2- tailed Student t test with Welch’s correction. **b** Histological pictures of kidney section from wild-type, *Ctns^-/-^*, *Gal3^-/-^*and *Ctns^-/-^ Gal3^-/-^* mice at 12-months-old stained with hematoxylin and eosin. Mononuclear infiltrates are indicated by arrows. **c** Kidney sections from wild-type, *Ctns^-/-^*, *Gal3^-/-^* and *Ctns^-/-^ Gal3^-/-^* mice at 8-9 months old were stained with anti-CD68 antibodies (green), phalloidin (white) and DAPI (blue). Scale bar: 50 µm. Quantification was then performed using Image J (wild-type: n=3; *Ctns^-/-^*, *Gal3^-/-^* and *Ctns^-/-^Gal3^-/-^*: n=4). *P* values were determined using one-way ANOVA followed by Tukey’s test. \*\**P* < 0.01.

We assessed the renal function by measuring creatinine, urea and phosphate levels in the serum and protein and phosphate in 24-hour urine collections. No significant difference was observed between WT and *Ctns^-/-^* mice at 8-9 months (data not shown). In contrast, at 12-15 months, *Ctns^-/-^* showed significantly higher serum creatinine and urea, compared to WT. *Ctns^-/-^ Gal3^-/-^* mice exhibited an improved renal function compared to *Ctns^-/-^* mice at 12-15 months of age, with lower serum creatinine (*P*<0.05) and urea (*P*<0.05) (Table 1).

**Table 1:** Serum and urine analyses for renal function of 12-15 months old mice

|  | Wild-type<br>(n=13) | *Ctns*<sup>-/-</sup><br>(n=16) | *Gal3*<sup>-/-</sup><br>(n=5) | *Ctns*<sup>-/-</sup> *Gal3*<sup>-/-</sup><br>(n=34) |
| --- | --- | --- | --- | --- |
| *Serum* |  |  |  |  |
| Creatinine (mg/dL) | 0.29 ± 0.02 | 0.41 ± 0.03<sup>a</sup> | 0.26 ± 0.02<sup>b</sup> | 0.31 ± 0.01<sup>b</sup> |
| Urea (mg/dL) | 48.95 ± 3.25 | 90.35 ± 3.34<sup>a</sup> | 52.68 ± 2.15<sup>b</sup> | 77.43 ± 2.49<sup>a,b,c</sup> |
| Phosphate (mg/dL) | 13.84 ± 1.07 | 12.87 ± 1.03 | 12.06 ± 0.28 | 12.03 ± 0.46 |
| *Urine* |  |  |  |  |
| Phosphate (μmol/24h) | 4.30 ± 0.62 | 5.56 ± 0.75 | 2.89 ± 0.81 | 6.06 ± 0.81 |
| Protein (mg/24h) | 12.36 ± 1.96 | 13.06 ± 2.04 | 14.69 ± 3.94 | 12.51 ± 1.42 |
| Volume (mL) | 0.85 ± 0.14 | 1.29 ± 0.14 | 0.89 ± 0.25 | 1.25 ± 0.12 |

*P* value was calculated by one-way ANOVA followed by Tukey's test.
<sup>a</sup> *P*<0.05 compared with wild-type mice
<sup>b</sup> *P*<0.05 compared with *Ctns*<sup>-/-</sup> mice
<sup>c</sup> *P*<0.05 compared with *Gal3*<sup>-/-</sup> mice

Histological studies of 8-9- and 12-15-month-old mice kidney sections stained with hematoxylin and eosin revealed the presence of severe anomalies in *Ctns^-/-^* mice kidney such as tubular atrophy, retracted glomeruli and mononuclear infiltrates. We performed blind analysis of kidney sections with scores ranging from 1 (preserved tissue structure) to 6 based on the percentage of extent of cortical damage. The average score for *Ctns^-/-^* mice was 2.64 ± 0.36 at 9 months of age (n=7) and 4.12 ± 0.37 at 12-15 months old (n=16). In the kidney of *Ctns^-/-^ Gal3^-/-^* mice at the same age, these anomalies were significantly less extensive than in the *Ctns^-/-^* kidney (Figure 4B); the averages scores were 1.62 ± 0.11 at 9 months (n=21; *P* <0.05, Mann-Whitney t test) and 3.04 ± 0.22 at 12-15 months (n=33; *P* <0.05, Mann-Whitney t test). Furthermore, a striking difference between *Ctns^-/-^* and *Ctns^-/-^ Gal3^-/-^* kidney sections was the presence of few mononuclear infiltrates in the *Ctns^-/-^ Gal3^-/-^* sections whereas heavy infiltrates were consistently observed in the *Ctns^-/-^* kidney (Figure 4B).

We characterized the mononuclear infiltrates observed by histology in 8-9-month-old *Ctns^-/-^* kidneys as macrophages/monocytes by immunostaining using anti-CD68 antibody (Figure 4C). The quantification revealed that significantly fewer macrophages/monocytes were observed in WT, *Gal3^-/-^* and *Ctns^-/-^ Gal3^-/-^* kidneys compared to *Ctns^-/-^* kidneys, confirming the data found by histology (Figure 4B). These data suggest that Gal3 is involved in the recruitment of macrophages/monocytes in cystinotic kidneys and that its absence ameliorates kidney disease in *Ctns^-/-^* mice. Hence, these data suggest that inflammation is responsible, at least in part, for kidney deterioration in *Ctns^-/-^* mice.

### *Ctns*-deficient kidneys exhibit Gal3 overexpression

We evaluated Gal3 expression in kidney explanted from 12-month-old WT and *Ctns^-/-^* mice by western blot analysis and found that Gal3 was significantly increased in *Ctns^-/-^* mice compared to WT (Figure 5A-B). To investigate further this increased expression of Gal3 in *Ctns^-/-^* kidneys, we quantified Gal3 expression at the transcript and protein levels in WT and *Ctns^-/-^* mice, and used as control kidneys from mice with chronic kidney disease (CKD) induced by standard subtotal two-step nephrectomy operation and from the sham-operated animals. *Gal3* transcripts were significantly increased in *Ctns^-/-^*and sham kidneys but much highly increased in CKD kidneys compared to WT mice (Figure 5C). In contrast, at the protein level Gal3 was detected by western blotting only in *Ctns^-/-^* kidneys (Figure 5D). Immunofluorescent staining of Gal3 in murine kidney at 8-9-month-old also demonstrated overexpression of Gal3 in the *Ctns^-/-^* kidney compared to WT (Figure 5E). Altogether, these data are consistent with decreased Gal3 degradation in absence of cystinosin as demonstrated *in vitro* (Figure 3A-B).

**Figure 5:**
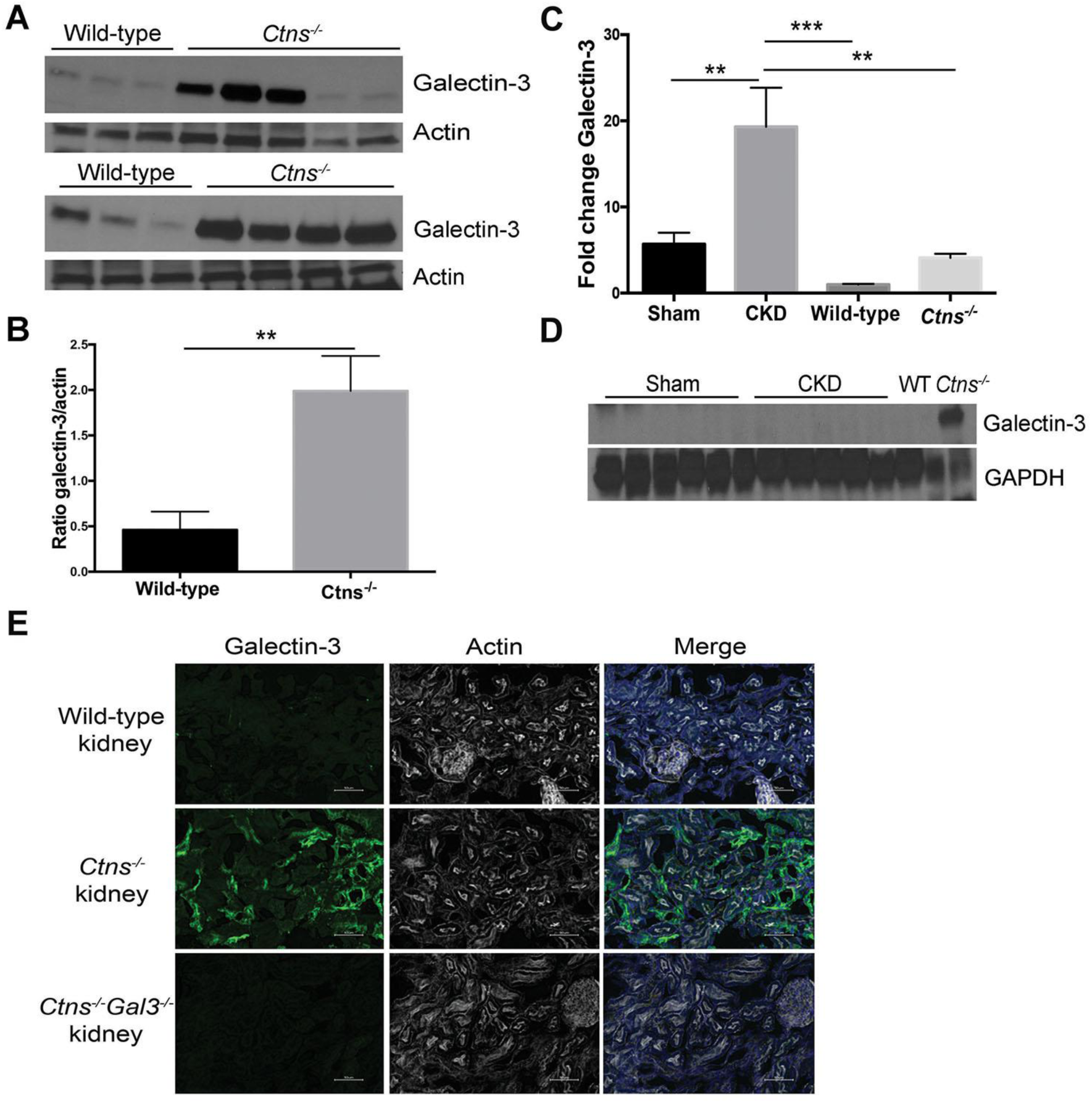
Gal3 expression is increased in the *Ctns^-/-^* mice. Gal3 expression in whole kidney homogenates of 12-month-old wild-type or *Ctns^-/-^* mice was evaluated by **a** western blot and **b** quantified using Image J (WT: n=6; *Ctns^-/-^:* n=9). *P* values were determined using unpaired 2-tailed Student t test with Welch’s correction. \*\**P*<0.01. mRNA and proteins were isolated from kidneys of sham-operated (n= 6) or 5/6 nephrectomized (CKD; n=6) animals and Gal3 expression was determined by droplet digital PCR **c** or western blot **d**. Protein lysate of kidney from WT and *Ctns^-/-^* mice at 12 months were used as negative and positive controls respectively for Gal3 expresion. *P* values were determined using one-way ANOVA followed by Tukey’s test. \*\**P*<0.01; \*\*\**P*<0.0001. **e** Wild-type, *Ctns^-/-^*and *Ctns^-/-^ Gal3^-/-^* kidneys were stained with anti-Gal3 antibodies (green), phalloidin (white) and DAPI (blue). Scale bar: 50 µm.

### Absence of cystinosin leads to increased serum MCP1 via Gal3 upregulation

Gal3 can regulate inflammation through different mechanisms (Henderson & Sethi, 2009). Thus, to gain further insights into the mechanism of Gal3-mediated inflammation, we investigated the expression of different pro- or anti-inflammatory cytokines in the serum of 12-15-month-old WT, *Ctns^-/-^, Gal3^-/-^* and *Ctns^-/-^ Gal3^-/-^* mice. Six cytokine levels were measured by mouse Cytometric Bead Array (CBA), MCP1, interferon-G (IFNG), tumor necrosis factor (TNF) and the following interleukins (IL) IL-10, IL-6 and IL-12p70. While no significant differences were observed for the other cytokines, MCP1 expression was significantly increased in *Ctns^-/-^* mice serum compared to WT and *Gal3^-/-^* mice and also to *Ctns^-/-^ Gal3^-/-^*mice, whose MCP1 serum levels were comparable to WT mice (Figure 6A). MCP1 is produced by different cell types but the major source of MCP1 are macrophages and monocytes (Deshmane, Kremlev et al., 2009). It is a chemokine known as a chemoattractant for monocytes and macrophages, which can regulate their migration and infiltration. In contrast, Gal3 expression was found similar in the serum of *Ctns^-/-^* mice (Mean ± SEM: 44.33 ± 9.81 ng/mL; n=5) and in WT mice (Mean ± SEM: 40.86 ± 6.41 ng/mL; n=7). Gal3 and MCP1 relationship was further investigated by co-immunoprecipitation using Gal3-GFP and MCP1-DsRed- or Gal3-GFP and CTNS- DsRed- (as a positive control) expressing 293 T cells. Interaction between Gal3 and MCP1 was observed in the bound fraction of the immunoprecipitation (Figure 6B), suggesting a direct induction of MCP1 by Gal3. Unbound fraction contains proteins that didn’t bind Gal3. Incubation with N-Acetyl-D-lactosamine (LacNAc), inhibitor of Gal3 CRD, showed that, in contrast to cystinosin interaction, the CRD is not involved in the interaction between Gal3 and MCP1 (Figure 6C). Cells transiently expressing Gal3-GFP alone, MCP-1 alone of Gal3-GFP and DsRed were used as controls (data not shown) and showed no non-specific interaction following immunoprecipitation.

**Figure 6:**
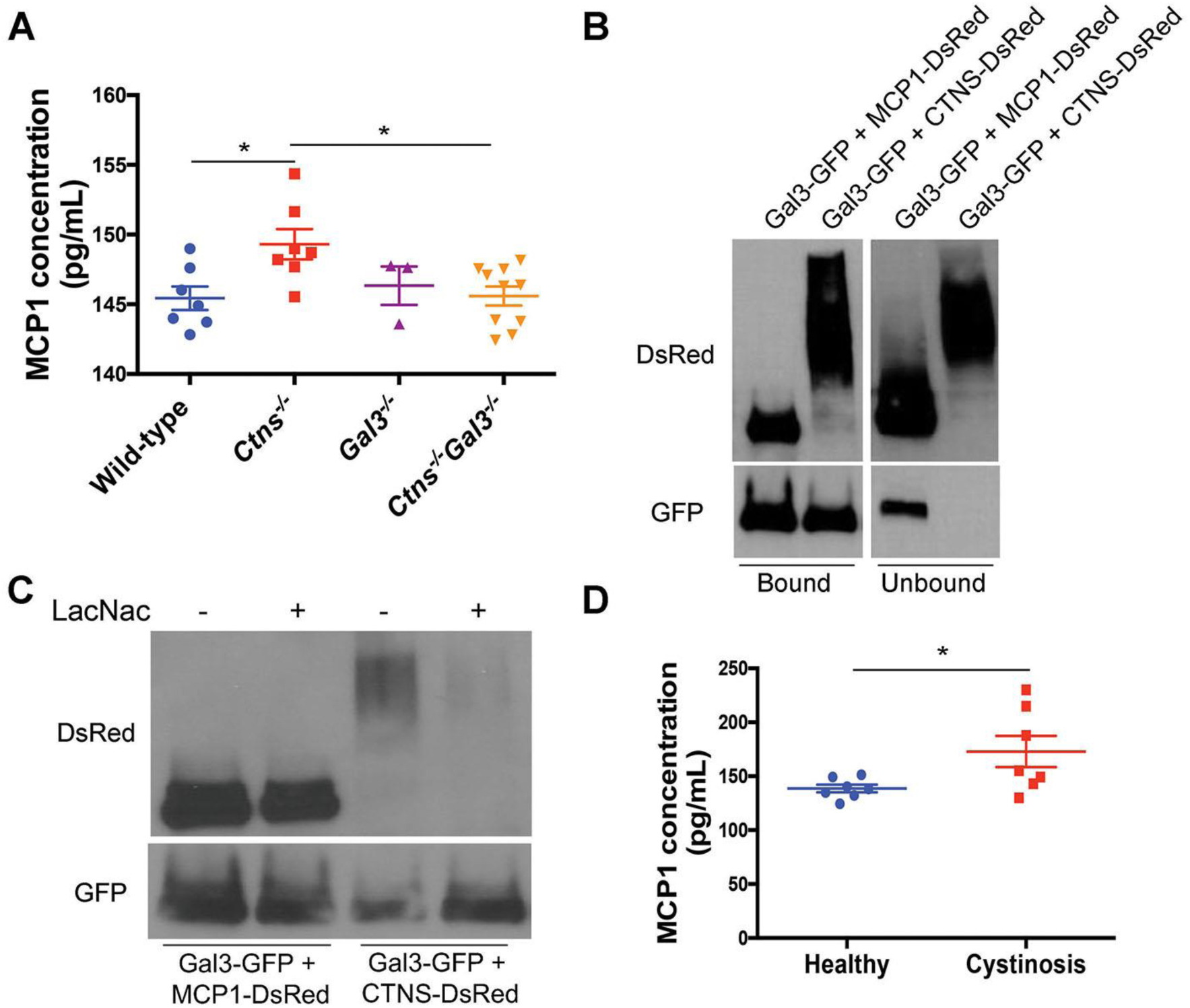
Gal3 interacts with MCP1, a macrophage/monocyte chemoattractant protein upregulated in cystinotic mice serum. **a** A Cytometric Beads Array has been used to determine MCP1 concentration in the serum of wild-type (n=7), *Ctns^-/-^* (n=7), *Gal3^-/-^*(n=3) and *Ctns^-/-^ Gal3^-/-^* mice (n=10). *P* values were determined using one-way ANOVA followed by Tukey’s test. \**P* < 0.05. **b** Lysates of 293T cells transfected with Gal3-GFP and MCP1-DsRed or Gal3-GFP and CTNS-DsRed (control) were immunoprecipitated with anti-GFP antibody and coimmunoprecipitated proteins were analyzed by western blotting. **c** The same lysates were treated or not with 5 mM of L-acetyl-D-lactosamine (LacNAc), a potent inhibitor of Gal3 carbohydrate interactions. Lysates were then immunoprecipitated with anti-GFP antibody and co-immunoprecipitated proteins were then analyzed by western blotting. **d** A Cytometric Beads Array directed against human MCP1 has been used to determine MCP1 expression in human serum samples from healthy donors (n=7) and cystinotic patients (n=7). *P* values were determined using unpaired 2-tailed Student’s t test. \**P* < 0.05.

To verify if these data are relevant in humans, we measured MCP1 levels in patients with cystinosis, who are not taking immunosuppressive therapy or anti-inflammatory medications, using a CBA directed against human MCP1 (Figure 6D). Serum MCP1 level was found significantly increased in patients with cystinosis (Mean ± SEM: 172.90 ± 14.53 pg/mL; n=7; mean age: 7.71) despite their young age, compared to healthy controls (Mean ± SEM: 138.6 ± 3.56 pg/mL, n=7; mean age: 37.6) (*P*<0.05). One additional patient with cystinosis, who is taking indomethacin, has a normal serum level of MCP-1, 114.93 pg/mL. Indomethacin can be used in cystinosis to reduce polyuria (Haycock, Al-Dahhan et al., 1982) but was also shown to inhibit galectin-3 and MCP1 expression (Dabelic, Flogel et al., 2005). Therefore, this patient is not included in the primary cohort.

## Discussion

Cystinosis is a lysosomal storage disorder characterized by accumulation of the amino acid cystine within cells. The kidneys are especially vulnerable to damage and most patient develop kidney failure by their second decade of life whereas cystine accumulate in all tissues. Thus, we believe cystine accumulation is not alone responsible of kidney degeneration. Herein, we describe a new role for cystinosin in the regulation of inflammation that we show as being involved in the progression of chronic renal disease in cystinosis.

The study of cystinosin molecular partners revealed a direct interaction with the member of the galectins family, Gal3. Gal3 is characterized by a CRD that binds beta-galactoside structures. The N-terminal domain of cystinosin is localized inside of the lysosomal lumen and contains 7 glycosylation sites (Nevo et al., 2017). Since interaction between cystinosin and Gal3 can be inhibited by addition of thiodigalactoside or N-acetyl-D-lactosamine, two substances that interacts with Gal3 CRD, interaction probably occurs through Gal3 CRD with cystinosin glycosylated moiety. While no signal peptide is present in Gal3, it is found in the nucleus, cytoplasm, at the cell surface or extracellularly and has a role in a variety of biological functions such as apoptosis, cell growth, cell adhesion and cell activation depending on its subcellular localization (Dumic et al., 2006). Here, we showed that Gal3 can also be localized within lysosomes and that its expression is decreased when cystinosin is overexpressed, strongly suggesting that cystinosin enhances Gal3 lysosomal localization leading to its degradation. Gal3 has previously been shown to be degraded through lysosomal-dependent proteolysis in absence of cAbl and Arg, two tyrosine kinases (Li et al., 2010). Here we showed that cystinosin and Gal3 colocalize at dynamic vesicular structures and are together mobilized in a spatiotemporal manner. Cystinosin has been previously associated to trafficking mechanisms of the lysosomal LAMP2A, the only known chaperone-mediated autophagy receptor, which is mislocalized in cystinosis (Napolitano et al., 2015b). Gal3 degradation has been previously shown to be mediated by CMA (Li et al., 2010). Confocal microscopy analysis confirmed the colocalization of Gal3 at Hsc70-positive structures in both *Ctns^-/-^*and WT cells, supporting the co-transportation of Gal-3 with Hsc70 for presentation at the lysosome and degradation by CMA. However, deficient localization of LAMP2A at the lysosome in cystinotic cells (Napolitano et al., 2015b) would help explain the decreased degradation of Gal3 in cystinosis. A possible interpretation of our results is that cystinosin regulates the trafficking and delivery of Gal3 to the CMA-active lysosomes for degradation. This novel pathway of cystinosin-dependent Gal3 lysosomal localization and degradation is supported *in vivo* as *Ctns*-deficient mice exhibit overexpression of Gal3 within the kidney. Moreover, while increased *Gal3* mRNA was limited in *Ctns^-/-^* kidneys compared to another model of CKD, large quantity of Gal3 protein was detected only in *Ctns^-/-^* kidneys. These data support that cystinosin is involved in the degradation of Gal3 protein, which can control overexpression of Gal3 mRNA in stress conditions.

Gal3 has several biological roles, including roles in acute and chronic inflammation by attracting monocytes and macrophages and by regulating alternative macrophage activation (MacKinnon, Farnworth et al., 2008, Sano, Hsu et al., 2000). In *Ctns^-/-^* mice, we observed heavy mononuclear infiltrates mainly in the kidney as previously described (Nevo et al., 2010, Yeagy, Harrison et al., 2011). We characterized the infiltrates as monocytes/macrophages. In contrast, kidneys from the double knock-out *Ctns^-/-^ Gal3^-/-^* mice exhibit very few monocyte/macrophage infiltrates, strongly suggesting that Gal3 is involved in inflammation in the kidneys of the *Ctns^-/-^*mice. In the context of repetitive tissue injury, Gal3 may trigger the transition to chronic inflammation and fibrosis (Henderson & Sethi, 2009). Consistent with these findings, our data demonstrate the involvement of Gal3 in the progression of the chronic kidney disease in cystinosis. Indeed, the double knockout *Ctns^-/-^ Gal3^-/-^* mice exhibit a better kidney function and structure compared to *Ctns^-/-^* mice. Henderson and colleagues showed that monocytes and macrophages express Gal3 and that its secretion leads to renal fibrosis (Henderson, Mackinnon et al., 2008). In their study, Gal3 deficient mice exhibit less collagen deposition and alpha smooth muscle actin expression in the kidney compared to WT mice after Unilateral Ureteric Obstruction (UUO). Moreover, absence of Gal3 protects against activation and accumulation of myofibroblasts known to promote fibrosis (Henderson et al., 2008, Henderson, Mackinnon et al., 2006).

O’Seaghdha et *al*. found a correlation between higher levels of Gal3 in plasma and development of chronic kidney disease (CKD) (O’Seaghdha, Hwang et al., 2013). However, in cystinosis, no difference was observed in the serum of *Ctns^-/-^*and WT mice. Measuring different cytokine levels in the serum using a microarray bead-based assay, we found a significant increase of MCP1, a monocyte/macrophage chemoattractant, in *Ctns^-/-^*mice while the *Ctns^-/-^ Gal3^-/-^* had level comparable to WT mice. MCP1 level was also found significantly increased in the serum of cystinosis patients compared to healthy donors. MCP1 can be released in the serum and form a gradient to attract monocytes and macrophages to the site of injury (Deshmane et al., 2009). In addition, we showed a direct interaction between Gal3 and MCP1, which is in contradiction with a previous study showing that Gal3 macrophages chemoattractant property was independent from MCP1 expression (Sano et al., 2000). An interaction between Gal3 and MCP1 was recently suggested by Gordon-Alonso *et al*. (Gordon-Alonso, Hirsch et al., 2017) but was not studied any further. Variation of Gal3 role in different disease states within the same organ such as the kidney has been described and implies the existence of different pathways in each context (Fernandes Bertocchi, Campanhole et al., 2008, Henderson et al., 2008, Nishiyama, Kobayashi et al., 2000, Okamura, Pasichnyk et al., 2011). In cystinosis, our data strongly suggest that the absence of cystinosin leads to the decrease of Gal3 degradation, which activates MCP1 blood mobilization, enhancing renal inflammation and progression of chronic kidney disease.

These findings open new perspectives in potential therapeutic targets that could limit or delay kidney degeneration in cystinosis patients. As such, indomethacin, a non-steroidal anti-inflammatory drug, is used to regulate polyuria and polydipsia in cystinosis (Haycock et al., 1982). A recent study on 307 European patients with cystinosis showed an improved renal outcome in patients treated with indomethacin (Emma, 2016). Indomethacin has been shown to inhibit Gal3 expression by activating the transcription factor PPARG that causes inhibition of activity of NFKB, a transcription factor implicated in Gal3 transcription (Dabelic et al., 2005). Moreover, de Borst et al, showed a decrease of MCP1 serum level in patients following a 4-week treatment with indomethacin (de Borst, Nauta et al., 2012). These findings are consistent with our data linking Gal3/MCP1 to the progression of the chronic kidney disease in cystinosis. Indomethacin is commonly used in patients with cystinosis in Europe but not in the US because some studies showed a potential nephrotoxicity of this drug (Lee, Cooper et al., 2007, Rocha, Michea et al., 2001). In light to our study, the use of indomethacin for cystinosis patients may be reconsidered.

## Methods

### Animal experiments

The C57BL/6 *Ctns^-/-^* mice were provided by Dr. Antignac (Inserm U1163, Paris, France) and bred continuously at University of California, San Diego. C57BL/6 galectin-3 null (*Gal3^-/-^*) mice were provided by Dr. Fu-Tong Liu (University of California, Davis) and Dr. Jerrold M. Olefsky (University of California, San Diego). *Ctns^-/-^* and *Gal3^-/-^* mice were backcrossed to obtain *Ctns^+/-^ Gal3^+/-^* mice. Mice with combined deficiency of *Ctns^-/-^* and *Gal3^-/-^* were generated by interbreeding of *Ctns^+/-^ Gal3^+/-^* animals. Using this breeding strategy, we also generated wild-type, *Ctns^-/-^*, *Gal3^-/-^* mice. All animal procedures complied with IACUC and NIH guidelines for the care and use of laboratory animals.

### Cell lines

MDCK type-II (ATCC) were grown in Dulbecco’s modified Eagle medium (DMEM) supplemented with 10% Fetal Calf Serum (FCS), 100 units/mL penicillin, 0.1mg/mL streptomycin and 2 mM L-glutamine. 293T cells were cultured in high glucose, L-glutamine supplemented DMEM with addition of 10% heat inactivated Fetal Bovine Serum (FBS), 100 units/mL penicillin and 0.1 mg/mL streptomycin. Mouse Embryonic Fibroblasts (MEF) were generated from newborn skin biopsies of wild-type and *Ctns^-/-^* C57BL/6 and were grown in high glucose, L-glutamine supplemented DMEM with addition of 10% heat inactivated Fetal Bovine Serum (FBS), 100 units/mL penicillin and 0.1 mg/mL streptomycin.

### Co-immunoprecipitation and mass spectrometry analysis

For investigation of protein-protein interactions, MDCK cells were transduced using the lentiviral pRRL.SIN.cPPT.PGK/WPRE vector backbone to stably express cystinosin-GFP (Dull, Zufferey et al., 1998). Cells were rinsed 3 times with cold phosphate buffered saline (PBS) and lysed in ice-cold lysis buffer (Tris 50mM, NaCl 150mM) containing 0,05% DDM (n-Dodecyl-β-D-maltoside, Fluka) and one tablet of Complete Protease Cocktail (Roche) per 50 ml. Lysates were cleared by 10 min centrifugation at 1000xg, 4°C and organelles were concentrated in supernatants by centrifugation at 100,000xg for 1 hour at 4°C. Protein concentration was determined by a BCA protein assay kit (Thermo Scientific), according to the manufacturer’s instructions. Co-immunoprecipitations were performed using µMACS GFP or Protein G Microbeads Isolation Kit (Miltenyi Biotec) for the isolation of GFP-tagged protein or Gal3, respectively. Briefly, 50 µl of MicroBeads conjugated to anti-GFP monoclonal antibody (Roche) were incubated with lysates containing 3 mg proteins at 4°C for 1 h, then applied onto µMACS separation columns. Columns were rinsed three times with 50mM Tris HCl, 300 mM NaCl, 1% Triton X-100, then three times with 50 mM Tris HCl, 150 mM NaCl, 1% Triton X-100 and finally three times with 20mM Tris HCl. Elution was performed with 50 µl of the kit elution buffer supplemented with 1% SDS. For Gal3 immunoprecipitation, 4 µg of anti-Gal3 antibodies (Cederlane) were incubated with 3 mg of lysates, overnight at 4°C and then 50 µl of Protein G MicroBeads were added and incubated for 1h at 4°C. Lysates were applied onto µMACS separation columns and protein isolation was performed as described above. Precipitated proteins were resolved by SDS-PAGE on 10% gel for mass spectrometry and immunoblotting. Coimmunoprecipitated protein samples prepared as above were treated and analysed as already described by mass spectrometry, LTQ Orbitrap (Andrzejewska, Nevo et al., 2015).

293T cells transiently transfected with Gal3-GFP and MCP1-DsRed or Gal3-GFP and CTNS-DsRed by the calcium phosphate co-precipitation method were rinsed with PBS and lysates in ice-cold lysis buffer containing 50mM Tris, 150 mM NaCl, 1% Triton X-100 and 1% proteinase inhibitor. 293T expressing Gal3-GFP alone, MCP-1 alone or Gal3-GFP and DsRed were used as controls. One mg of proteins from the lysates was incubated with 4 µg of anti-GFP antibody (Roche) overnight at 4°C. Samples were incubated with 50 µL of Protein G Microbeads for 1h at 4°C before separation on a magnetic board. Beads were washed 3 times with ice-cold buffer, resuspended in lysis buffer containing 1X of LDS, 1X of reducing agent and 1X of proteinase inhibitor and heated at 55°C for 10 minutes to separate the proteins from the beads. Proteins were resolved by SDS-PAGE and Gal3-binding MCP-1 was detected using anti-DsRed antibody (Clontech).

### Subcellular Fractionation

Livers were obtained from C57BL/6 mice. The preparation was performed on eight mouse livers essentially as described previously (Wattiaux, Wattiaux-De Coninck et al., 1978). Briefly, fractionation of subcellular organelles by differential centrifugation produced nucleus and heavy mitochondrial (NM), light mitochondrial (L), and microsomal and soluble (PS) fractions. The L fraction was subjected to isopycnic centrifugation on a discontinuous Nycodenz density gradient. Conditions of the gradient were essentially the same as described in the original publication (Wattiaux et al., 1978), except that Nycodenz® was used instead of metrizamide. The following density layers were successively loaded on top of the L fraction: 1.16 (7 ml), 1.145 (6 ml), 1.135 (7 ml), and 1.10 (7 ml). Centrifugation was performed at 83,000xg for 2 h 30 min in an SW28 Beckman rotor. Fraction 2, the interface between layers of respective densities, 1.10 and 1.135 g/mL, was recovered as the “lysosome-enriched” fraction (LEF). This LEF was diluted in 0.25 M sucrose, pelleted by centrifugation (30,000xg, 4°C, 10 min), and homogenized in 300 µl of sucrose. Protein concentration was determined by a BCA protein assay kit (Thermo Scientific), according to the manufacturer’s instructions.

### Gal3-inhibitor and Proteinase K treatments

Lysates from cystinosin-GFP MDCK cell line were incubated in the presence or in the absence of 5 mM thiodigalactoside (TDG), a specific competitive inhibitor of Gal3. Lysates from 293T transiently expressing Gal3-GFP and CTNS-DsRed or Gal3-GFP and MCP1-DsRed were incubated with 5mM of N-Acetyl-D-Lactosamine (LacNac), an inhibitor of Gal3 CRD. After 30 min of incubation with TDG or LacNac, at 4°C with constant shaking, lysates were incubated with 50 µl anti-GFP MicroBeads and immunoprecipitation was performed as mentioned above. Precipitated proteins were resolved by SDS-PAGE on 10% gel.

Lysates from cystinosin-GFP MDCK cell lines and lysosome-enriched fractions (LEF) from mouse liver were first incubated with or without 2% Triton X-100 for 10 min at 4°C. Then lysates and LEF were incubated 30 min with or without 2.5 µg/ml and 25 µg/ml proteinase K, respectively. Enzyme treatment was stopped by incubation with 1 mM phenylmethylsulfonyl fluoride 5 min at 4°C. Proteins were resolved by SDS-PAGE on 10% gel.

### Immunofluorescence analysis

MDCK cells were rinsed with PBS, fixed with 4% formaldehyde in PBS at room temperature for 20 min followed by treatment with 50 mM NH_4_Cl for 10 min. Cells were incubated with primary antibodies, anti-Lamp-2 antibody (Acris) and anti-Gal3 antibody (Cederlane) diluted 1:100 in PBS containing 0.075% saponin and 0.1% BSA for 2 hours at room temperature and then with secondary antibody Alexa Fluor 555 (Life Technologies) diluted 1:200 in PBS containing 0.075% saponin and 0.1% BSA for 1 h at room temperature. Confocal images were taken using a ZEISS LSM 700 scanning laser confocal microscope (Carl Zeiss MicroImaging GmbH, Jena, Germany).

293T cells were transfected using calcium chloride to obtain cells transiently expressing Gal3-GFP alone, Gal3-GFP and DsRed or Gal3-GFP and cystinosin-DsRed. These cells were cultured on a coverslip and, 48h after transfection, the coverslip was mounted on a slide using mounting medium. The cells were then imaged using a Keyence BZ-X700 (Keyence).

Kidneys were fixed in 4% paraformaldehyde for 30 minutes, equilibrated in 20% sucrose overnight and frozen in Tissue-Tek Optimal Cutting Temperature (Sakura Finetek) buffer at -80°C. Sections of 10 μm were blocked with 1% bovine serum albumin, 5% donkey serum in phosphate buffer saline. The blocking buffer was then used to dilute primary and secondary antibodies. Kidney sections were stained overnight at 4°C with anti-CD68 antibody (Biolegend) or anti-Gal3 antibody (Biolegend), dilution 1:100 followed by Alexa Fluor 488 donkey anti-rat IgG (Life Technologies), 1h at room temperature. Samples were finally stained with Alexa Fluor 647 phalloidin (Life Technologies), dilution 1:2000 and DAPI (Thermofisher) for 30 minutes at room temperature. Images were acquired using a Keyence BZ-X700 fluorescent microscope (Keyence). To quantify CD68 expression, nine images were acquired for each kidney sections and stitched together to obtain one image using BZ-X700 software. Images were then analyzed using ImageJ 1.48v software (National Institutes of Health).

MEFs were transfected with Gal3-GFP with the Neon transfection system (Thermo Fisher), following the manufacturer’s instructions. The cells were then stained using anti-Hsc70 antibody (Enzo) and imaged as described previously (Zhang, Johnson et al., 2017)

### Total Internal Reflection Fluorescence (TIRF) microscopy

WT and *Ctns^-/-^* MEF were transfected with vectors for the expression of Gal3-GFP and CTNS-DsRed with the Neon transfection system (Thermo Fisher), following the manufacturer’s instructions. The cells were seeded on 4-chamber 35 mm-borosilicate bottom dish (In Vitro Scientific). Two days post-transfection, MEFs were analyzed by TIRF microscopy. TIRF microscopy experiments were performed using a 100X 1.45 numerical aperture TIRF objective (Nikon) on a Nikon TE2000U microscope custom-modified with a TIRF illumination module as described (Johnson, Napolitano et al., 2013). Laser illumination (488 and 568 nm laser lines) was adjusted to impinge on the coverslip at an angle to yield a calculated evanescent field depth of 300-400 nm (Pseudo-TIRFM), a technique that increases the deepness of analysis yet maintaining the high signal to background ratio of traditional TIRF. Images were acquired on a 16-bit, cooled charge-coupled device camera (Hamamatsu) controlled through NIS-Elements software. For live experiments, the images were recorded using 300–500 ms exposure time depending on the fluorescence intensity of the sample. The images were analyzed using Image*J*.

### Gal3 expression analysis *in vitro*

293T cells transfected with Gal3-GFP, Gal3-GFP and DsRed or Gal3-GFP and CTNS-DsRed using calcium chloride coprecipitation method were lysed in RIPA buffer containing a proteinase inhibitor cocktail. Protein concentration was determined using Pierce BCA Protein Assay Kit (Thermo Fisher Scientific). 10 μg of proteins were separated using a 4-15% Mini-protean TGX gel (Biorad) and transferred onto polyvinylidene difluoride membrane. After blocking, the membrane was incubated with anti-Gal3 antibody (Abcam), dilution 1:500 or anti-GAPDH (Cell signaling), dilution 1:2000. Immunoreactive bands were detected using horseradish peroxidase (HRP)-conjugated antibody (Abcam), dilution 1/5000 and ECL Western Blotting Substrate (Thermo Scientific).

### Renal function

Just prior to sacrifice, 24-hour urine was collected in metabolic cages. Serum was obtained by retro orbital bleeds. Serum and urine phosphate levels, serum creatinine and urea were estimated using Quantichrom Phosphate Assay Kit, Quantichrom Creatinine Kit and Quantichrom Urea Assay Kit (Bioassay Systems). Protein levels in urine were measured using Pierce BCA Protein Assay Kit (Rockford, IL).

### Histology

At time of sacrifice, kidneys were collected, fixed in formalin and embedded in paraffin. Sections were stained with hematoxylin and eosin and reviewed in a blinded fashion by Dr. Marie-Claire Gubler. Blinded scoring was done in a range from 1 to 6 based on the extent of cortical damage estimated in % terms: 1 (0-10% damaged kidney), 2 (10 to 30%), 3 (30 to 50%) 4 (50 to 70%), 5 (70 to 90%) and 6 (>90%).

### Cystine content measurement

Tissue cystine measurement was performed as previously described (Yeagy et al., 2011). Briefly, explanted tissues were grounded in N-ethylmaleimide (Fluka Biochemika) using Precellys 24 (Bertin). Proteins were precipitated using 15% Sulfosalicylic acid (SSA; Fluka Biochemika) resuspended in NaOH 0.1N and measured using the Pierce BCA protein assay kit (Thermo Fisher Scientific). The cystine-containing supernatants were sent to the UCSD Biochemical Genetics laboratory to be analyzed by mass spectrometry (LC-ESI-MS/MS).

### Standard Nephrectomy and Sham operation

CKD in mice was induced by standard subtotal two-stage nephrectomy operation as described previously (Cheung, Yu et al., 2005, Cheung, Kuo et al., 2007). Briefly, the CKD in mice was established by a two-phased procedure of 5/6 nephrectomy. After anesthesia, mice were put into left lateral position, shaved, disinfected with 75 % ethanol.

Locating the kidney under rib ridge and cutting a about 2 cm opening on skin, then cut the muscles; use tweezers to pull up the kidney out of surrounding fat, peel off renal cell membrane from the lower end (to avoid injury to adrenal gland); cut off about 1/3 renal tissue near both ends but keep parts around the renal hilum to avoid disruption of blood supply; wrap remaining kidney twice with a thick line and stretch 10 min with hemostat. When there was no further bleeding, the opening was sutured layer by layer and wiped clean with cotton swab. A week later the right renal hilum was ligated with thick lines to make kidney necrosis. The sham group of mice were undergone the same procedure but without cutting any kidney tissue.

### Gal3 expression analysis *in vivo*

Explanted kidneys were homogenized in RLT buffer complemented with β-mercaptoethanol or RIPA buffer complemented with protease inhibitor cocktail using Precellys 24 (Bertin). RNA was isolated using AllPrep DNA/RNA Mini Kit (Qiagen) and 200 ng of RNA was reverse transcribed using iScript cDNA Synthesis kit (Biorad). Gal3-specific droplet digital PCR was performed using 2 μl of cDNA, ddPCR Supermix for probes (No dUTP) (BioRad), Gal3 primers and probe (Applied biosystem, 4331182) and 18S primers and probe (Applied biosystem, 4319413E). The expression level of Gal3 gene, expressed as ratio values, was calculated by dividing the concentration of the gene of interest by an endogenous control (18S).

For protein isolation, homogenized samples were centrifuged and supernatants were collected. Protein concentration was determined using Pierce BCA Protein Assay Kit (Thermo Fisher Scientific). 25 μg of proteins were separated using a 4-15% Mini-protean TGX gel (Biorad) and transferred onto polyvinylidene difluoride membrane. After blocking, the membrane was incubated with anti-Gal3 antibody (Abcam), dilution 1:500. Immunoreactive bands were detected using horseradish peroxidase (HRP)-conjugated antibody (Abcam), dilution 1/5000 and ECL Western Blotting Substrate (Thermo Scientific).

An ELISA (ab203369, Abcam) was used to detect Gal3 expression in mice serum. Briefly, serum samples (dilution 1:1000) were incubated with a cocktail of capture antibody and reporter conjugated detector antibody for one hour, on a plate shaker at room temperature. After washes, Tetramethylbenzidine (TMB) substrate was added to the wells and the plate was incubated for 10 minutes, shaking at room temperature. The reaction was stopped and the plate was read using a spectrophotometer at an OD of 450 nm.

### Cytokine level quantification in serum

To measure cytokine levels in mouse serum we used a mouse inflammation kit Cytometric Bead Array from BD Biosciences. The assay was performed according to manufacturer’s instruction. Briefly, 25 μL of serum (dilution 1:2) were stained with the mixture of mouse cytokine capture bead suspension and the phycoerythin (PE) detection reagent. Samples were incubated for 2 hours at room temperature and then washed and analyzed using BD Accuri C6 apparatus. Mouse cytokine standards provided with the kit were diluted appropriately to generate the standard curve. To measure the concentration of MCP1 in human serum, we used a Cytometric Bead Array human MCP1 Flex Set with a Human Soluble Protein Master Buffer Kit (BD Biosciences). According to manufacturer’s instructions, human serum samples (dilution 1:10) were incubated for one hour with Human MCP1 Capture Bead at room temperature. The PE detection reagent was then added and samples were incubated for another 2 hours. Beads were then washed and analyzed with a BD Accuri C6 (BD Bioscience). Human MCP-1 standards were appropriately diluted to obtain a standard curve.

### Statistical analysis

Values are expressed as mean ± SEM. The significance of the results was assessed by unpaired 2-tailed t-test or unpaired 2-tailed t-test with Welch’s correction. Group comparisons of 3 conditions or more were made with parametric analyses of variance, followed by Tukey’s multiple comparisons test for pairwise comparisons. Histological scores were compared using Mann-Whitney test. All analyses were performed using PRISM 6 software (GraphPad). A *P* value less than 0.05 was considered significant.

### Study approval

All experiments involving mice were conducted in compliance with approved Institutional Animal Use Committee protocols at Laboratory of Hereditary Kidney Diseases (Paris, France) and University of California, San Diego (San Diego, California, USA). Use of human tissue in this study was approved by the University of California, San Diego Human Research Protections Program.

## Acknowledgments

We acknowledge Dr. Fu-Tong Liu (University of California, Davis) and Dr. Jerrold M. Olefsky (University of California, San Diego) for providing the Gal3^-/-^ mice. Thank you to Lou Devanneaux and Nichole Flerchinger for their technical help. We acknowledge Christopher Alfonso from BD Bioscience for providing the mouse inflammation cytometric bead array and for his help with the interpretation of the results. This work was supported by the National Institute of Health (NIH) RO1-DK090058 and R01-DK110162, the Cystinosis Research Foundation and the California Institute of Regenerative Medicine (CIRM, CLIN-09230). T.L. is supported by a graduate fellowship from the Vaincre les Maladies Lysosomales. AB and JZ are supported by a fellowship from the Cystinosis Research Foundation. UCSD Neuroscience Microscopy Shared Facility was funded by the Grant P30-NS047101.

## Author contributions

TL, NN, CR, SDC, CW, RM, TM, CA and SC developed methodology. TL, RM, NN, JZ, AB and CG performed experiments. TL, NN, JZ, SDC, MCG, CA and SC analyzed the data. Resources were provided by SDC, RM, CA and SC. TL, SDC, CA and SC wrote the manuscript. TL, SDC, RM, TM, CA and SC reviewed and edited the manuscript.

## The paper explained

### Problem

Cystinosis is a rare lysosomal storage disorder and the kidney pathology associated with this disease is not fully understood. Treatment with cysteamine, which allows the exit of the cystine out of the lysosome, doesn’t prevent end stage renal failure and patients still need kidney transplantation. New roles for cystinosin, other than lysosomal cystine transporter, have been recently identified, and these findings may allow to the discovery of new therapeutic targets.

### Results

Using unbiased interaction screening, a novel interaction was found between cystinosin, the protein implicated in cystinosis, and a member of the galectin family, galectin-3 (Gal3). We showed that cystinosin enhanced lysosomal localization of this protein and its degradation. These data were confirmed *in vivo* since Gal3 was found increased in the kidney of cystinotic mice. Creation of the mouse model deficient for both Gal3 and cystinosin revealed a better renal function and structure, and a decrease of pro-inflammatory cells infiltrates in the *Ctns^-/-^ Gal3^-/-^*mice. By looking at pro-inflammatory cytokines, we found an increase expression of Monocyte Chemoattractant Protein-1 (MCP-1) in the serum of cystinotic mice and patients, and demonstrated direct interaction between Gal3 and MCP-1. This cytokine is implicated in the recruitment of monocytes and macrophages in inflammation revealing a new inflammatory pathway in cystinosis.

### Impact

Here, we demonstrate a mechanism in which cystinosin is involved in the regulation of inflammation via Gal3 interaction, that can impact the progression of the renal chronic disease in cystinosis. These findings suggest that treatments inhibiting or decreasing inflammation in cystinosis could lead to a delayed progression of the disease. Indomethacin, a nonsteroidal anti-inflammatory used in patients with cystinosis to decrease water intake, was shown to inhibit Gal3 and MCP-1 expression. Due to potential nephrotoxicity of this drug, it is not commonly prescribed to patients in the United States. However, it is commonly used in Europe and a recent study performed in Italy showed that indomethacin was actually beneficial for the kidney pathology in cystinotic patients. The present study could lead to the reconsideration of the use of this drug in cystinosis.

